# Structure-based discovery of a novel small-molecule inhibitor of TEAD palmitoylation with anticancer activity

**DOI:** 10.1101/2022.06.09.495565

**Authors:** Artem Gridnev, Subhajit Maity, Jyoti R. Misra

**Affiliations:** Department of Biological Sciences, University of Texas at Dallas, Richardson, TX, United States

**Keywords:** YAP, TEAD, Hippo signaling, cancer, Small-molecule inhibitor, Virtual ligand screening

## Abstract

The paralogous oncogenic transcriptional coactivators YAP and TAZ are the distal effectors of the Hippo signaling pathway, which play a critical role in cell proliferation, survival and cell fate specification. The core kinase cascade of the Hippo signaling pathway phosphorylates YAP/TAZ and restrict its activity by promoting cytoplasmic sequestration and degradation. Unphosphorylated YAP/TAZ translocates into the nucleus and associates with the TEA domain (TEAD1-4) transcription factors to regulate their target genes. They are frequently misregulated in most human cancers, where they contribute to multiple aspects of tumorigenesis including growth, metabolism, metastasis and chemo/immunotherapy resistance. Thus, they provide a critical point for therapeutic intervention. However, due to their intrinsically disordered structure, they are challenging to target directly. Since YAP/TAZ exerts oncogenic activity by associating with the TEAD1-4 transcription factors, to regulate target gene expression, YAP activity can be controlled indirectly by regulating TEAD1-4. Interestingly, TEADs undergo autopalmitoylation, which is essential for their stability and function. The palmitic acid occupies a hydrophobic pocket in TEAD1-4, and small-molecule inhibitors that bind to this site can render them unstable, and allosterically inhibit their interaction with YAP/TAZ and chromatin. Here we combined structure-based virtual ligand screening with biochemical and cell biological studies and identified a novel small-molecule that inhibits TEAD palmitoylation and stability. Further, it affects YAP-TEAD interaction and inhibits YAP transcriptional activity and impairs proliferation, colony formation and migration of breast and ovarian cancer cells.

## 1 INTRODUCTION

The Hippo signaling pathway is a conserved signaling network that plays a critical role in cell proliferation, survival, differentiation and tissue homeostasis (1, 2). The pathway consists of a core kinase cascade that negatively regulates the paralogous oncogenic transcriptional coactivators, Yes Associated Protein (YAP) and Transcriptional Activator with PDZ-binding motif (TAZ). The kinase cascade consists of the serine threonine kinases MST1/2 and Large Tumor Suppressor 1/2 (LATS1/2), and their obligate adapters SAV and MOB1A/B respectively, where MST1/2 phosphorylates and activates LATS1/2, which in turn phosphorylates YAP/TAZ. Lats1/2 can also be phosphorylated and activated by MAP4K1-7 and Tao1/3 kinases (3, 4). Phosphorylated YAP/TAZ gets sequestered in the cytoplasm, and gets ubiquitinated and degraded. Under low Hippo pathway activity, Hypo/unphosphorylated YAP/TAZ translocates into the nucleus, where it associates with various transcription factors such as TEAD1-4, SMAD3 and RUNX (5, 6). Of these transcription factors, TEAD1-4 mediate the regulation of majority of the YAP-target genes, which encode various cytokines and matricellular proteins that promote cell proliferation and inhibit apoptosis.

YAP/TAZ has emerged as a central player in many cancers including breast, colorectal, liver, lung, pancreas, thyroid and sarcomas (7, 8). Despite these observations, point mutations within the Hippo pathway components are relatively rare in most cancers. Most cancers harbor mutations in the upstream regulators that promote elevated expression and nuclear localization of YAP/TAZ. One of the key upstream regulator of YAP is Merlin, which is encoded by the NF2 gene. Merlin plays an important role in recruiting the core components of the Hippo signaling pathway to the plasma membrane, facilitating activation of the LATS1/2 kinase (9). Germline loss-of-function mutations or deletion of NF2 results in neurofibromatosis type 2, which causes bilateral vestibular schwannomas (10). Somatic mutations of NF2 are also observed in spontaneous schwannomas, meningiomas, mesothelioma and renal cell cancer. A comprehensive study of 32 different cancers revealed that malignant mesothelioma has the highest frequency of NF2 mutations (11, 12). Similarly, a large proportion of meningiomas harbor mutations in NF2 (13). Furthermore, 6% of non-small-cell lung cancer exhibit YAP amplification, while 29% of them show TAZ amplification (14). YAP is also frequently amplified in head and neck cancers. 90% of epithelioid hemangioendothelioma (EHE), a rare vascular sarcoma, harbor TAZ-CMTA1 fusion while 10% of EHE have YAP-TFE3 fusion (15-18). In addition to these, oncogenic mutations that activate growth promoting signaling pathways such as EGFR, RAS-MAPK, PI3K and Wnt signaling also promote higher YAP/TAZ expression.

Overexpressed YAP/TAZ undergoes phase separation at the super enhancers, and promotes sustained expression of the target genes (19-21). Genetic analyses revealed that YAP contributes to multiple aspects of cancer development, including growth, survival, metabolism and metastasis. It also plays an important role in cancer fibroblast proliferation and ECM deposition. They also promote cancer stem cell fate maintenance and chemotherapy resistance (22, 23). Furthermore, YAP promotes expression of the chemokine CXCL5, which results in the recruitment of myeloid cells that suppress T-cells (24). In regulatory T-cells (Tregs) YAP supports FOXP3 expression via activin signaling and Treg function. Accordingly, YAP deficiency results in dysfunctional regulatory T cells (Tregs), which are no longer able to suppress antitumor immunity (25). YAP upregulates PD-L1 expression in the cancer cells, and by this mechanism directly mediates evasion of cytotoxic T-cell immune responses (19, 22, 26-28). Thus, YAP/TAZ regulates multiple aspects of tumorigenesis, and provides a critical control point for therapeutic intervention for cancer treatment. However, it is challenging to directly inhibit YAP/TAZ, since it is an intrinsically disordered protein. Similarly, it is difficult to interfere YAP-TEAD1-4 interaction with small-molecules, as the interaction interface is very broad, shallow, and exposed to solvent, although a few molecules that bind to the interface-2 have been reported to achieve this. Several groups have developed linear and cyclic peptides that bind to TEAD and interfere with YAP/TAZ-TEAD interaction (29, 30). However, these molecules exhibit poor efficacy or cell permeability, which has limited their use. Therefore, recent efforts are aimed at alternate approaches to inhibit YAP activity indirectly, by disrupting TEAD function.

TEAD transcription factors undergo covalent modification with palmitic acid at a conserved cysteine residue, and are regulated by the APT2 and ABHD17 depalmitoylases (31, 32). The palmitic acid occupies a central hydrophobic pocket in these proteins and regulates their stability and activity. Inhibition of TEAD palmitoylation can render these proteins unstable, and allosterically interferes with their interaction with YAP/TAZ (33-35). Further, a small molecule inhibitor that binds to the palmitic acid binding pocket is known to exert a dominant negative effect on TEAD binding to the chromatin, and converts TEAD from a transcriptional activator to a transcriptional repressor (34). Interestingly, the central hydrophobic pocket is highly druggable, and more importantly, structural alignment has revealed that 75% of the residues are identical and the 25% of the residues are 75% similar across all the 4 TEAD isoforms (8). Thus, it is possible to develop pan-TEAD inhibitors that bind to this region. Several investigational compounds have been reported to bind to this site and inhibit YAP activity (29, 30). However, due to high failure rates during clinical trials and potential development of resistance, there is a continuing urgent need for developing novel chemotype-based potent TEAD inhibitors.

Recent developments in structure-based virtual ligand screening (VLS) allows one to conduct *in silico* screening of large libraries of small molecules to predict the ones, which have high probability of binding to a given site in a target protein (36-38). This significantly reduces the time, labor and cost to identify potential hit compounds. Here we report isolation of a novel small molecule that inhibits YAP transcriptional activity. Using structure-based VLS combined with cell biological and biochemical analyses we identified JM7 as a potent inhibitor of TEAD palmitoylation. It negatively regulates TEAD stability and YAP target gene expression. Furthermore, we show that this compound inhibits, proliferation, colony formation and migration of breast and ovarian cancer cell lines that exhibit high YAP/TAZ activity.

## 2 METHODS

### 2.1 Molecular Docking

The X-ray crystal structure of TEAD2 was retrieved from the protein Data Bank (PDB ID: 6UYC) and was prepared in the protein preparation wizard of Schrodinger. Missing side chains were added using Prime implemented in Maestro. Hydrogens were added and bond orders were assigned. The resulting structure was protonated at pH 7.0 using PROPKA. The ligands were retrieved from PubChem, prepared using LigPrep wizard in Maestro and protonated at pH 7.0 using Epik. Subsequently, energy minimization was performed using OPLS5 force field within 0.3A° root mean square deviation. Docking was performed using the Ligand docking workflow in Glide using Standard Precision mode (Glide SP). The top 25% of the compounds was subjected to Xtra precision (XP) docking and MM-GBSA. The compounds with the lowest dg-Bind score were chosen for testing how they affect YAP transcriptional reporter.

### 2.2 Molecular Biology

pCMX-GAL4-TEAD1 (Addgene #33108), pCMX-GAL4-TEAD2 (Addgene #33107), pCMX-GAL4-TEAD3 (Addgene #33106), pCMX-GAL4-TEAD4 (Addgene #33105), pRK5-Myc-TEAD4 (Addgene #24638), GST-YAP2 (Addgene #24637), pCMV-FLAG-YAP-5SA/S94A (Addgene #33103), and pCDNA3-HA-TAZ (Addgene #32839) were obtained from Addgene. The human codon-optimized TEAD3 coding sequence with N-terminal Myc tag was synthesized as a gBlock (Integrated DNA Technologies). Full length TEAD1 and TEAD2 coding sequences were amplified from pCMX-GAL4-TEAD1 and pCMX-GAL4-TEAD2 and cloned into the EcoRI site of pCDNA3.1 by Gibson assembly using the NEBuilder® HiFi DNA Assembly Kit (New England Biolabs Cat. #E2621). The TEAD3 gBlock fragment was similarly cloned into the EcoRI site of pCDNA3.1 by Gibson assembly. pCDNA3.1-TAZ-S89A was generated by creating overlapping fragments to mutate the target residue and ligated by Gibson assembly into the EcoRI site of pCDNA3.1. SmBiT-YAP, SmBiT-TAZ and LgBiT-TEAD1 were synthesized as gBlock fragments and cloned into the EcoRI site of pCDNA3.1 by Gibson assembly. The LgBiT-TEAD1 subsequently used as template to amplify the LgBit fragment of Nanoluc; TEAD2-YBD was amplified from pCMX-GAL4-TEAD2, TEAD3-YBD from pDONR221-TEAD3, and TEAD4-YBD from pRK5-Myc-TEAD4. pCDNA3.1-LgBiT-TEAD2, pCDNA3.1-LgBiT-TEAD3 and pCDNA3.1-LgBiT-TEAD4 were then constructed by Gibson assembly of their respective fragments into the EcoRI site of pCDNA3.1. All the primers used for cloning in this study are listed in Supplementary Table-1.

### 2.3 Cell Culture and Transfections

HEK293, MDA-MB-231, and OVCAR-8 cells were cultured in Dulbecco’s Modified Eagle’s Medium (DMEM, Corning Cat. #10-013-CV) supplemented with 10% heat-inactivated fetal bovine serum (HI FBS, Gibco™ Cat. #10438026) and 1% antibiotic/antimycotic (Gibco™). All cells were cultured in a humidified incubator at 37°C with 5% CO_2_. HEK293 cells were transiently transfected using Lipofectamine 3000 (Invitrogen), following the manufacturer’s protocol.

### 2.4 TEAD Transcriptional Reporter Assay

8XGTIIC lentiviral particles were purchased from BPS Bioscience, Inc. and used to transduce HEK293 cells. Stable clones were established by selection with 2.5 μg/mL puromycin until distinct colonies formed. Colonies were established and stable clones were verified for YAP-dependent induction of luciferase expression by overexpressing YAP-5SA/S94A or TAZ-S89. For dose response assays, HEK293-8XGTIIC cells were transfected with either YAP-5SA/S94A or TAZ-S89A. One day after transfection, cells were seeded in triplicate in 96-well plates and treated with either vehicle control or JM7 (1, 2 and 5 μM) for 24 hr. A duplicate of each plate was seeded to allow for normalization to ATP levels by Cell Titer Glow. Luciferase assays were performed using a one-step luciferase assay kit (Promega), and ATP-based viability with Cell-Titer Glo 2.0 (Promega), following the manufacturer’s protocol. Luminescence data was recorded on a Promega Glo-Max Navigator instrument.

### 2.5 RNA Extraction and RT-qPCR

MDA-MB-231 and OVCAR-8 cells were seeded at ∼50% confluence in 6 well dishes and allowed to attach overnight. Cells were then treated with DMSO or JM7 (2 μM and 5 μM) for 24 hr. Total RNA was extracted with TRIzol reagent (Invitrogen), following the manufacturer’s protocol. 1 μg of total RNA was reverse transcribed to cDNA using the iScript™ gDNA Clear cDNA synthesis kit (Bio-RAD). qPCR was performed with Applied Biosystems Fast SYBR Green master mix on an Applied Biosystems QuantStudio 6 Flex system. Target genes were normalized to GAPDH mRNA levels and relative fold changes calculated as 2^-ΔΔCt^.

### 2.6 Immunoprecipitation

Following transfection with the required plasmids and/or treatments, cells were trypsinized and collected in centrifuge tubes. Pellets were washed once with PBS, then lysed on ice for 30 minutes with PBS containing 0.5% IGEPAL-CA630 (Sigma-Aldrich) and cOmplete™ EDTA-free protease inhibitor cocktail. Following sonication and centrifugation at 4°C for 20 min at maximum speed, supernatants were collected and added to tubes containing 20 μL Pierce Protein A Agarose bead slurry (Thermo Scientific) and pre-cleared for 1 hr at 4°C. The pre-cleared supernatants were then added to tubes containing 50 μL Pierce anti-c-Myc agarose, EZview Red anti-FLAG/HA/Myc or V5-agarose affinity resins and incubated for 2 hr at 4°C. Immunoprecipitated protein was washed four times with PBS containing 0.5% IGEPAL-CA630 before proceeding to downstream assays such as palmitoylation assays or Western blotting.

### 2.7 Western Blotting

Samples were treated with Laemmli buffer, boiled at 98°C for 5 minutes, then centrifuged at maximum speed for 2 min following which, the samples were resolved by SDS-PAGE and transferred to nitrocellulose membranes using a Bio-RAD Trans-Blot Turbo semi-dry transfer system. After blocking with protein-free blocking buffer (Li-COR), blots were stained at 4°C overnight with anti-Myc and anti-actin primary antibodies. Next day, the blots were washed 4 times with PBST (PBS+1% Tween 20) and incubated with IR Dye 680 or IR Dye 780 conjugated secondary antibodies, or IR Dye 680 conjugated streptavidin (for palmitoylated fractions) for 1 hour at room temperature. After washing 4 times with PBST, the blots were scanned on a Li-COR Odyssey CLx system.

### 2.8 TEAD Palmitoylation Assay

HEK-293 cells transfected with Myc-TEAD1, Myc-TEAD2, Myc-TEAD3, or Myc-TEAD4 expression plasmids were treated with DMSO or 2 μM JM7 in conjunction with 100 μM alkyne palmitate for 24 hr. Myc-tagged protein was immunoprecipitated and palmitic acid was conjugated to biotin azide by click chemistry, where the precipitated TEAD proteins were incubated with 100 μL of reaction buffer containing 0.1mM biotin-azide, 1mM tris (2-carboxyethyl)phosphine (TCEP), 0.2mM tris (3-hydroxypropyltriazolylmethyl) amine (THPTA) and 1 mM CuSO_4_. The reaction was carried out at 20°C for 1 hr with shaking at 1200 rpm and terminated by washing three times with PBS containing 0.5% IGEPAL-CA630. Subsequently, the samples were subjected to Western Blotting using IR Dye 680 conjugated streptavidin to detect the biotinylated palmitic acid.

### 2.9 Immunofluorescence

MDA-MB-231 and OVCAR-8 cells were seeded in 4-well chamber slides and allowed to attach overnight. Cells were then treated with DMSO or JM7 (2 μM) for 24 hr. After treatment, the media was removed, cells were washed with PBS, then fixed for 20 min with 4% PFA, permeabilized for 10 min with 0.5% Triton X-100 in PBS, then blocked for 30 min with 1% donkey serum and 0.1% Triton X-100 in PBS, followed by staining with anti-YAP (D8H1X, Cell Signaling Technology #14074) and anti-pan-TEAD (D3F7L, Cell Signaling Technology #13295) at 4°C overnight, then with appropriate Alexa Fluor 647-conjugated secondary antibodies for 1 hr at RT and counterstained nuclei with Hoechst for 5 min at RT. Slides were then overlaid with coverslip with Vectashield mounting medium and sealed with nail polish. Images were captured on a Leica TCS SP8 confocal laser scanning microscope and processed in Volocity.

### 2.10 MTT assay

MTT assay was performed using the MTT assay kit (Abcam) following manufacturer’s protocol. Briefly, equal number of MDA-MB-231 or OVCAR-8 cells were plated in triplicate and were treated with DMSO or 2uM JM7 for 48 hours, following which they were incubated with the MTT reagent for 1 hours. The formazan crystals that formed were dissolved in the solubilization buffer and the absorbance was measured at 590nm wave length.

### 2.11 Colony formation assay

Equal number of MDA-MB-231 or OVCAR-8 cells were plated in triplicate and were treated with DMSO or 2uM JM7 for two weeks, following which they were fixed with 4% PFA and stained with 0.5% crystal violet dissolved in methanol. The images were acquired using Gelcount (Oxford Optronix) mammalian-cell colony, spheroid and organoid counter, and the colony areas were quantified using FIJI.

### 2.12 Wound Healing/Scratch Assay

MDA-MB-231 or OVCAR-8 cells were grown at full confluence and a scratch was made using a pipette tip. The detached cells were gently washed off with PBS and cells were incubated with media containing either DMSO or 2uM JM7. Images of the scratch areas were taken at 0, 19 and 24 hours following drug treatment and analyzed using FIJI.

### 2.13 NanoBiT Complementation Assay

HEK-293 cells were seeded at a density of 2 × 10^5^ cells/mL in a 24 well dish and transfected with 250 ng of SmBiT-YAP, SmBiT-TAZ, LgBiT-TEAD1/2/3/4 alone or in combination. One day after transfection, cells were seeded in triplicate in 96-well plates and treated with either DMSO or JM7 (2 μM) for 24 hr. Nanoluc assays were performed using the Nano-Glo Luciferase assay kit, following the manufacturer’s protocol. Luminescence was measured in a Promega Glo-Max Navigator instrument.

### 2.14 Statistical Analysis

Results were recorded and sorted in Microsoft Excel and all statistical analyses were carried out using GraphPad Prism (San Diego, CA). Histograms show the mean plus Standard Error of Mean (SEM). For pairwise comparisons we used two-tailed Student’s t test for parametric distributions and a Mann-Whitney test for non-parametric distributions.

## 3 RESULTS

### 3.1 Molecular Docking Based Identification of JM7

To identify potential small-molecule inhibitors that bind to the TEAD central pocket, where the palmitic acid normally resides, we conducted a structure-based VLS. To achieve this, we used the crystal structure of TEAD2 with PDB ID 6UYC and prepared it for docking using the Protein Preparation Wizard in Glide. This structure has a resolution of 1.66 angstroms and is co-crystalized with a ligand that binds with 229 nanomolar affinity (34). We generated a library of about 600,000 small molecules from PubChem and prepared the ligands for docking using Ligprep Wizard in Glide. After performing standard precision (SP) docking the top 2% compounds were subjected to extra precision (XP) docking in Glide. Finally, Molecular Mechanics with Generalized Born and Surface Area solvation (MM/GBSA) was performed to allow 5 A° movement of the protein structure. The top 50 compounds with highest free energy of binding (dG-bind score) were purchased and examined if they inhibit YAP transcriptional activity.

To screen the compounds for inhibiting YAP-transcriptional activity, we generated a stable HEK-293 cell line that enables to monitor YAP-TAZ/TEAD transcriptional activity, by infecting with a lentivirus carrying 8 tandem copies of the TEAD binding site from the SV40 enhancer (8XGTIIC), upstream of a Firefly luciferase (Fluc). To normalize this reporter activity, we either measured the ATP levels by CellTiterGlow™ or transfected with a plasmid expressing Renilla luciferase from the HSV thymidine kinase promoter (Rluc). Given that the basal Fluc activity of the 8XGTIIC reporter is low, we transfected these cells with plasmids encoding unphosphorylatable YAP (YAP5SA) or TAZ (TAZS89A). Expression of these mutant proteins dramatically activated the Fluc expression, as expected, without affecting Rluc expression (Figure S1. A, B). We then treated these cells with 10uM of the compounds to examine if they inhibited the reporter activation. From this screen we identified a compound JM7 (Figure 1. A), which inhibited the YAP transcriptional reporter, without causing any apparent effect on cell viability. It is predicted to bind to the TEAD central pocket mostly through hydrophobic interactions, hydrogen bonding with Ser345 and pi-pi stacking with Tyr426 (Figure 1. B-D).

**Figure 1.**
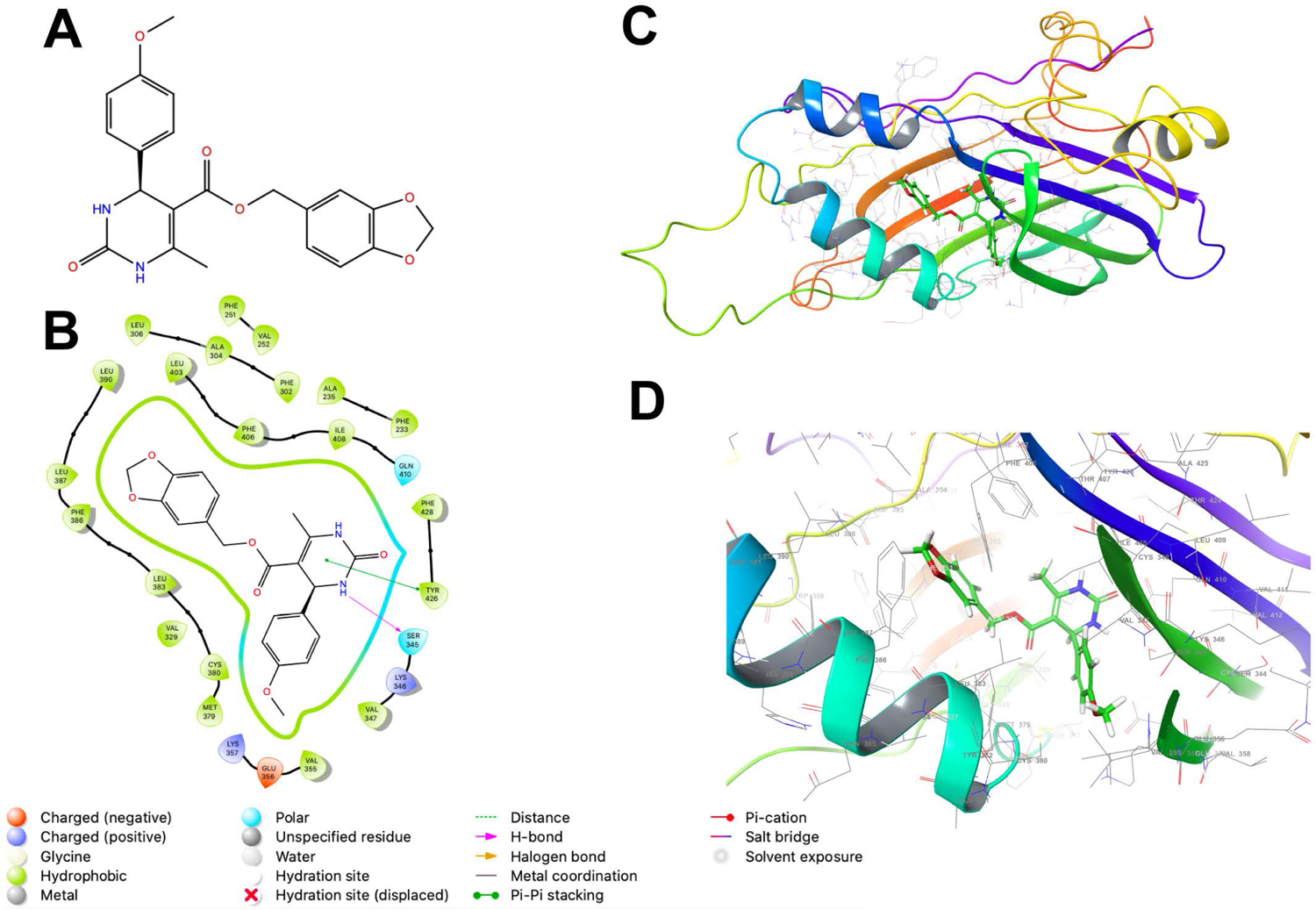
JM7 is predicted to bind to the TEAD cetral pocket. (A) Chemical structure of JM7. (B,D) Structure of TEAD2 (PDB ID: 6UYC) with predicted binding pose of JM7 and enlarged view (D). (C) Schematic showing the ligand interaction diagram for JM7. Please also see Figure S1.

To examine if JM7 inhibits YAP/TAZ activity in a dose-dependent manner, we treated HEK293-8XGTIIC cells expressing YAP 5SA or TAZ S89A with DMSO alone or 1, 2 or 5 uM JM7 overnight and examined how they affected YAP transcriptional activity. We observed that JM7 caused a dose-dependent decrease in Fluc expression induced by YAP 5SA (Figure 2. A-A”) and TAZ S89A (Figure 2 B-B”). In parallel we performed CellTiter Glow™ assay, which measures the general cell viability. We observed no significant decrease in CellTiter Glow activity in cells treated with different doses of JM7 compared to the vehicle treated controls cells (Figure 2. A’, B’). To determine the IC50 value of JM7, we transfected HEK293 cells expressing the 8XGTIIC-Fluc reporter with plasmids encoding Rluc and YAP5SA, following which we treated them with logarithmic concentrations of JM7 and determined the concentration at which it inhibits Fluc expression by 50% (Figure 2. C). In parallel, we also measured the Rluc expression (Figure 2.D). We observed that JM7 caused a 50% inhibition of Fluc expression at 972 nanommolar concentration. Together, these experiments indicate that JM7 inhibits YAP transcriptional activity in a dose-dependent manner with an IC50 value of 972nanomolar.

**Figure 2.**
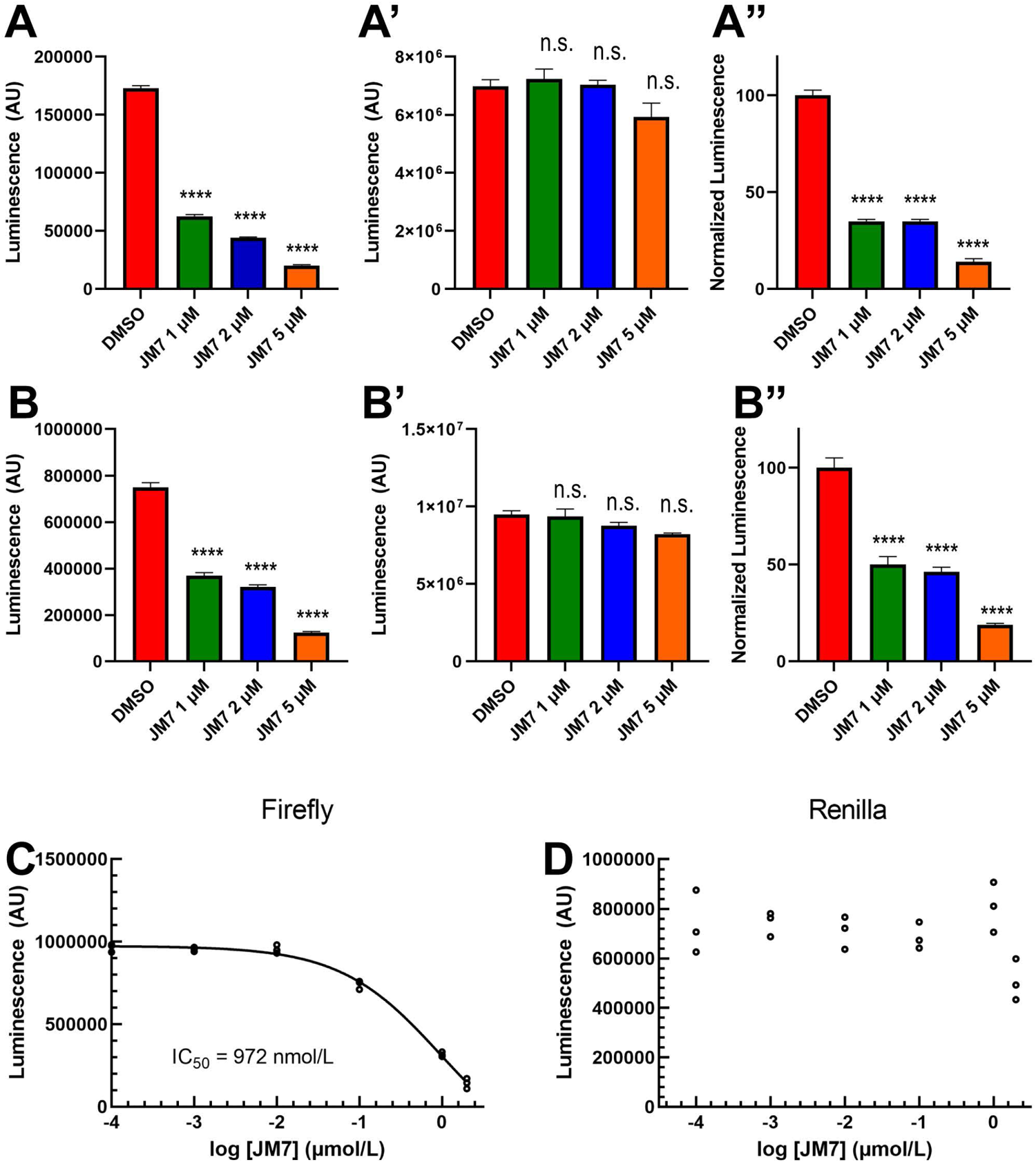
JM7 inhibits YAP/TAZ transcriptional activity. (A-B”) Luciferase reporter activity (A, B), Cell Titer Glow activity (A’, B”) and the normalized luciferase reporter activity (ratio of Luciferase reporter activity to Cell Titer Glow activity normalized to DMSO treated cells) (A”, B”) in HEK293 cells carrying 8xGTIIC-Luc and expressing YAP5SA (A-A”) or TAZ S89A (B-B”), treated with DMSO or indicated doses of JM7 showing dose dependent inhibition of reporter activity without apparent effect on cell titer glow activity. C, D) Normalized Fire fly luciferase or Renilla luciferase activity in HEK293 cells carrying 8xGTIIC-FLuc, pCMV-RLuc and YAP5SA treated with different doses of JM7 showing IC50 value of 972nM. ****=p<0.0001, ns= not significant. Error bars indicate Standard Error of Mean (SEM).

**Figure 3.**
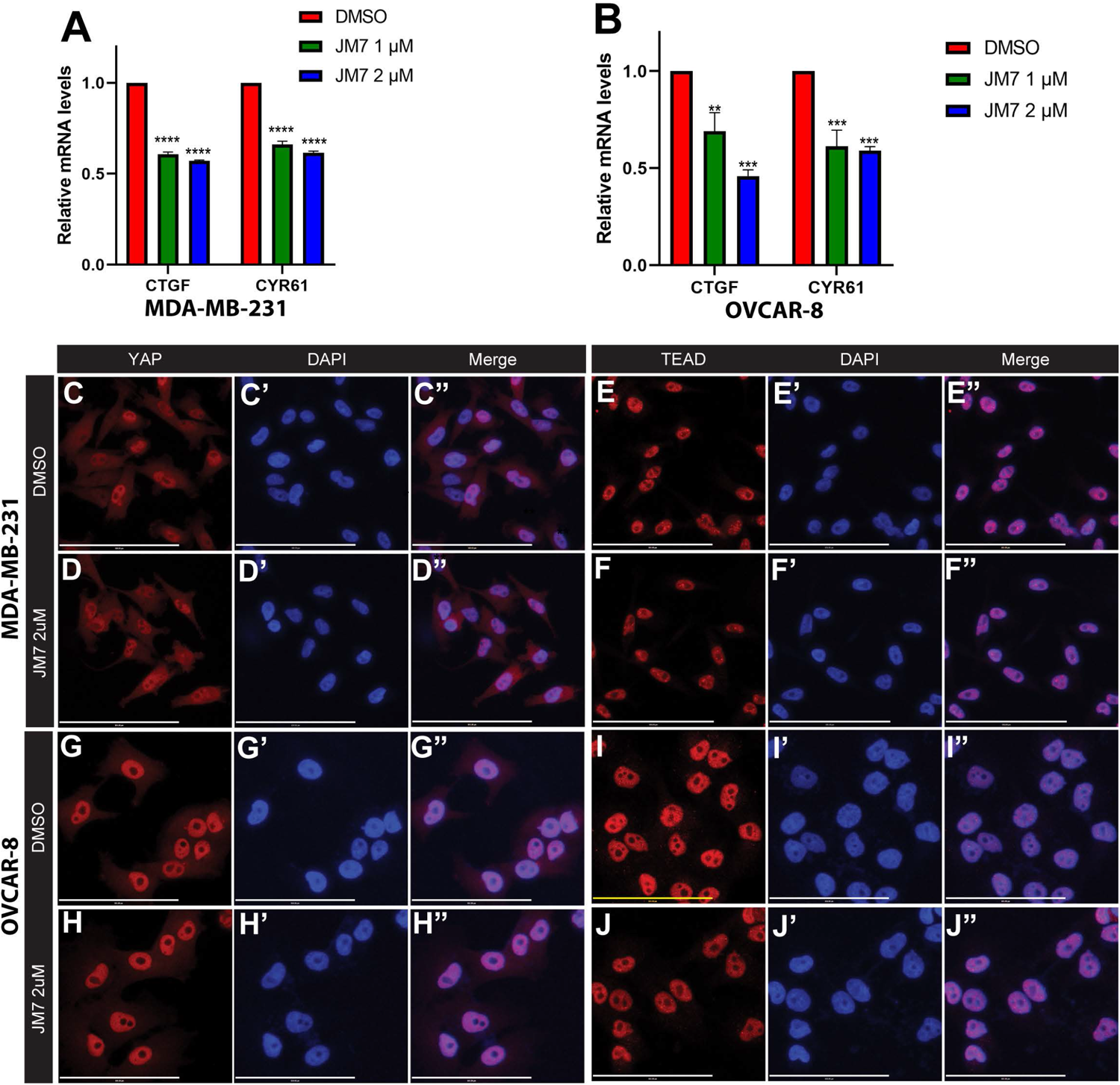
JM7 inhibits YAP target genes in breast and ovarian cancer cells and does not affect YAP or TEAD localization. (A-B) Histograms showing relative expression of CTGF and CYR61 mRNA levels in MDA-MB-231 (A) and OVCAR-8 (B) cells treated with DMSO or indicated doses of JM7, showing a dose dependent inhibition of these YAP target genes. (C-J”) MDA-MB-231 (C-F”) and OVCAR-8 (G-J”) cells were treated with DMSO or 2 micromolar JM7 and stained with TEAD or YAP antibody and Hoechst to stain the nuclei showing that JM7 does not seem to affect YAP or TEAD nuclear localization. ** p< 0.05; *** p<*0.005;**** p<0.0001. Scale bar=100um. Error bars indicate Standard Error of Mean (SEM).

### 3.2 JM7 Inhibits YAP Target Gene Expression

Encouraged by the effect of JM7 on YAP/TAZ transcriptional reporter activity, we then sought if it affects the expression of YAP/TAZ target genes. *CTGF* and *CYR61* are two of the well characterized YAP target genes, and are overexpressed in cancer cells that exhibit increased YAP activity. MDA-MB-231 is a triple negative breast cancer cell line that harbors a deletion in the NF2 gene, and therefore, exhibits increased YAP activity. Similarly, OVCAR-8 ovarian cells overexpress TEAD4. Therefore, we treated MDA-MB-231 and OVCAR-8 cells with DMSO alone or 1, 2 or 5uM JM7 and examined the CTGF and CYR61 transcript levels by quantitative reverse transcription polymerase chain reaction (qRT-PCR). GAPDH transcript levels were used for normalizing the expression levels. We observed that JM7 significantly downregulated the expression of both CTGF and CYR61 transcript levels in a dose dependent manner, in both MDA-MB-231 and OVCAR-8 cells (Figure2. A, B). Together, these results indicate that JM7 not only inhibits YAP transcriptional reporter activity, but also inhibits YAP target gene expression.

### 3.3 JM7 Does Not Affect Nuclear Localization of YAP and TEAD

YAP and TEAD activity is primarily controlled by their nuclear localization, and an inhibitor can block their activity, potentially, by affecting their levels and preventing their nuclear translocation. Therefore, we examined if JM7 affects YAP/TEAD localization in MDA-MB-231 and OVCAR-8 cells. We treated these cells with DMSO alone or 2uM JM7 and stained with α-YAP and α-TEAD antibodies and counterstained the nuclei with Hoechst. In the DMSO treated cells, as expected, TEAD primarily localized to the nucleus and YAP localized to both cytoplasm and the nucleus. Similarly, JM7 treated cells displayed similar localization of YAP and TEAD as the DMSO treated cells (Figure 2. C-J”). These results suggest that JM7 does not affect YAP/TEAD nuclear localization to inhibit their transcriptional activity.

### 3.4 JM7 Inhibits TEAD Palmitoylation and Protein Stability

Since JM7 was predicted to bind to the central hydrophobic pocket that is normally occupied by the palmitic acid, we sought to examine if it affects TEAD palmitoylation. To address this, we transfected HEK-293 cells with plasmids expressing Myc epitope tagged TEAD1-4 and treated them with alkyne palmitic acid along with DMSO or 2 micromolar JM7 for 24 hours. Myc-TEAD1-4 was subsequently immunoprecipitated using anti-Myc affinity resins and the alkyne palmitate was covalently conjugated with azide-biotin using click chemistry. Subsequently, palmitoylation was detected by Western blotting with fluorescently labeled Streptavidin. Because JM7 treatment destabilizes TEAD1-4, to equalize total Myc-TEAD1-4, less amount of the DMSO treated We observed that 2 micromolar JM7 significantly inhibits palmitoylation of all four TEAD isoforms, albeit to different extent (Fig 4.A-D). The effect was more pronounced in case of TEAD-1, TEAD-2 and TEAD-4 than in TEAD-3. Together, these experiments indicate that JM7 inhibits TEA palmitoylation.

**Figure 4.**
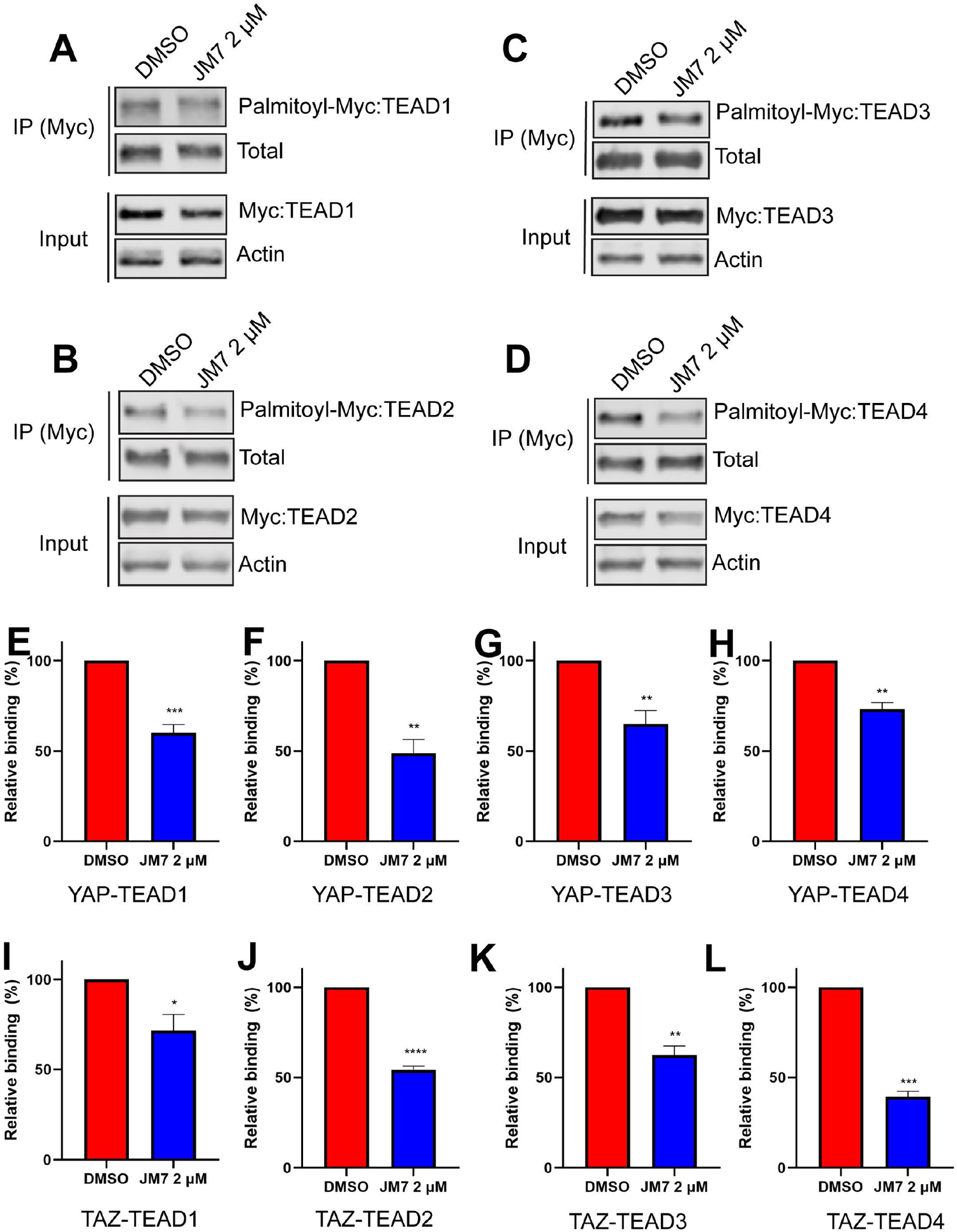
JM7 inhibits TEAD palmitoylation, impairs TEAD stability and inhibits YAP/TAZ-TEAD interaction. Western blots showing decreased palmitoylation (in the IP lanes) and degradation (in the input lanes) of TEAD1 (A) TEAD2 (B) TEAD3 (C) and TEAD4 (D) in lysates of HEK293 cells expressing Myc tagged TEAD1-4 and treated with DMSO or 2 micromolar JM7,. Actin is used as a control for equal loading and transfer of the input samples. In the IP samples, less amount of DMSO treated sample was loaded to match the amount of total TEAD in the JM7 treated samples. HEK293 expressing Myc tagged TEAD1-4 were treated with DMSO or 2 micromolar JM7 along with alkyne palmitic acid. The Myc-TEAD was immunoprecipitated and the alkyne palmitic acid was covalently conjugated with azide-biotin by click chemistry and the biotin was detected by blotting with fluorescently conjugated streptavidin.(E-L) Histograms showing relative NanoLuc activity in DMSO and JM7 treated HEK293 cells expressing SmBit-Yap (E-H) or Smbit-TAZ (I-L) together with TEAD1(E, I), TEAD2(F, J), TEAD3 (G, K) and TEAD4 (H, L). Error bars indicate Standard Error of Mean. ** p< 0.05; *** p<*0.005; **** p<0.0001.

**Figure 5.**
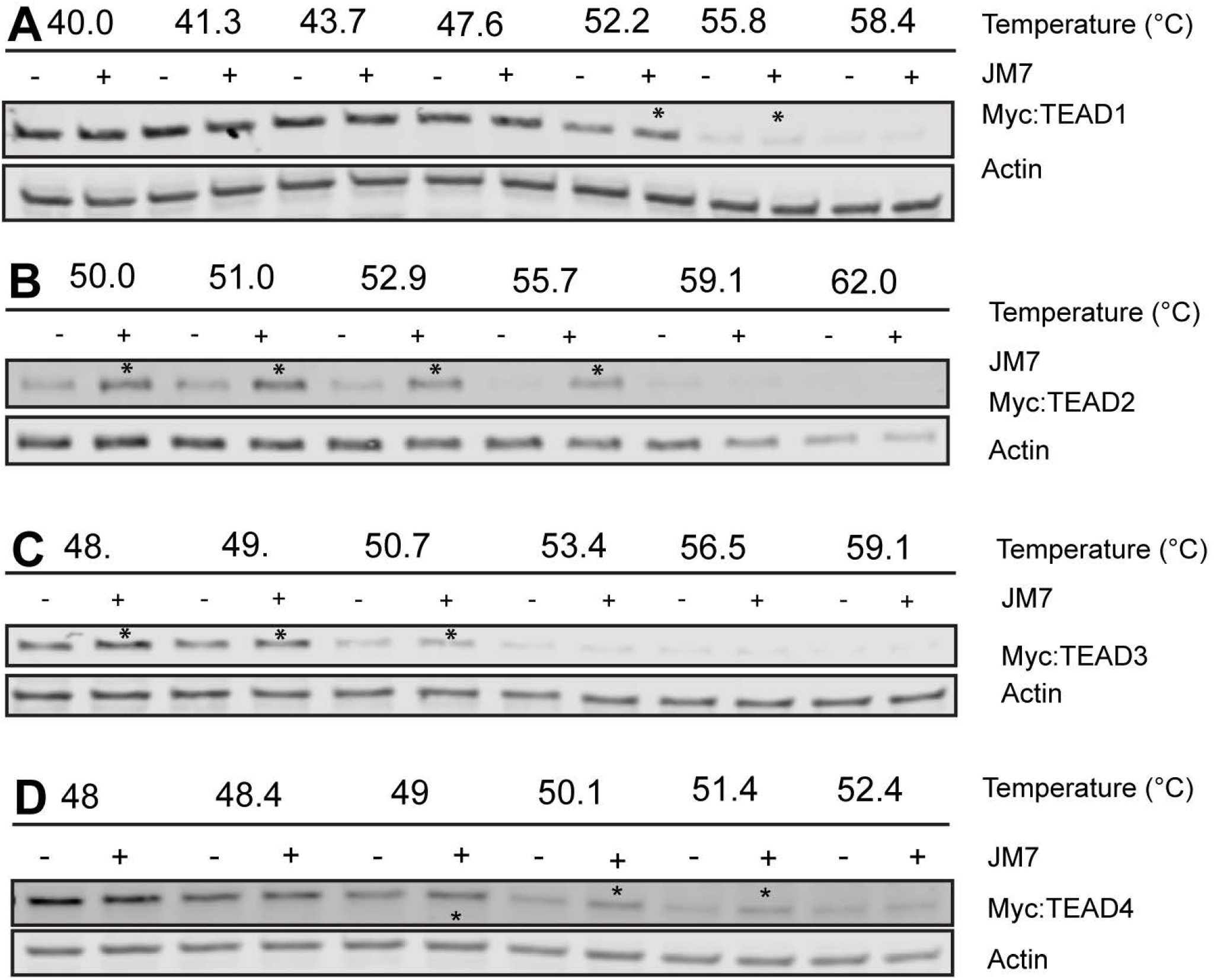
JM7 engages with TEAD1-3 in cells. (A-D) Representative Western blots showing amount of Myc-TEAD1-4 present in the supernatant following Cellular Thermal Shift Assay (CETSA). HEK293 cells expressing Myc-TEAD1-4 were treated with DMSO or 10 micromolar JM7 for 4 hours to avoid TEAD degradation, and CETSA was performed in triplicate. Asterisks indicate stabilization of JM7-bound TEAD.

Palmitoylation is critical for TEAD stability, and inhibition of this modification with certain small-molecule inhibitors destabilize these proteins. However, one small-molecule TEAD inhibitor that binds to the palmitic acid binding pocket was reported to stabilize TEAD, but dominant negatively interfere with its association with chromatin (34). In order to examine how JM7 affects TEAD stability, we transfected HEK293 cells with plasmids encoding Myc-TEAD1-4 and treated them with either DMSO or 2uM JM7 for 24 hours and examined the levels of Myc-TEAD by Western blotting, using anti-Myc antibody. We observed that JM7 affects the stability of all four TEAD isoforms to different extent. The destabilizing effect was more pronounced in case of TEAD-1 and TEAD-4, while the effect on TEAD2 and TEAD3 stability was less apparent (Figure 4, A-D). The weaker effect on TEAD3 stability could be due to less efficient inhibition of palmitoylation, while in case of TEAD2 could result from higher stability of this isoform.

### 3.5 JM7 Inhibits YAP-TEAD Interaction

Small molecule inhibitors that bind to the TEAD central pocket and inhibit their palmitoylation, can allosterically modulate its interaction with YAP/TAZ (33-35). Therefore, we examined if JM7 interferes with YAP/TAZ-TEAD interaction. To test this, we used the NanoBit complementation assay (39). In this assay, an engineered small bright Luciferase called Nanoluc is split into two fragments, SmBit and LgBit that do not interact with each other to reconstitute the enzymatic activity. However, when these fragments are fused to YAP and TEAD respectively, YAP-TEAD interaction brings these fragments in close proximity and reconstitutes the Nanoluc activity. Any compound that inhibits YAP-TEAD interaction would therefore cause a decrease in Nanoluc activity. This system was originally developed for YAP and TEAD-1. Now we have generated the fusion proteins to test interaction of both YAP and TAZ with all four TEAD isoforms. As expected, cells expressing SmBit-YAP, SmBit-TAZ or LgBit-TEAD1-4 alone had very low basal luminescence (Figure S2 A-D). However, cells expressing SmBit-YAP or SmBit-TAZ together with the different isoforms of LgBit-TEAD exhibit very high luminescence. When these cells were treated with either DMSO or 2 micromolar JM7, we observed that JM7 treatment significantly decreased Nanoluc activity (Figure 4. E-L), suggesting that JM7 interferes with YAP/TAZ-TEAD interaction.

### 3.6 JM7 Directly Engages with TEAD1-4 in Cells

Given that JM7 inhibits TEAD transcriptional activity and inhibits their palmitoylation, we wanted to examine if JM7 directly binds to the TEADs. To test this, we performed cellular thermal shift assay (CETSA), which is based on the principle that ligand bound proteins are resistant to thermal denaturation compared to their unbound counterparts (40). We expressed Myc-tagged TEAD1-4 in HEK293 cells and treated them with either DMSO or JM7, after which we subjected the cells to thermal denaturation at a gradient of increasing temperatures. The cells were subsequently lysed by freeze thawing and the denatured proteins were separated from the non-denatured proteins by centrifugation. The supernatant containing non-denatured proteins was examined for Myc-TEAD by Western blotting using anti-Myc antibody. We observed that while at high temperature, both DMSO and JM7 treated Myc-TEAD1-4 gets denatured, at lower temperatures, JM7 treated samples contain higher amount of Myc-TEAD, compared to DMSO treated samples, indicating that JM7 directly binds to TEAD1-4 and renders them resistant to thermal denaturation. Further, the thermal shift was more pronounced in case of TEAD2 and TEAD-3 than TEAD1. Together, these experiments suggest that JM7 directly engages TEAD1-4 in cells.

### 3.7 JM7 inhibits proliferation, colony formation and migration in breast and ovarian cancer cells

Since JM7 binds to TEAD and inhibits its palmitoylation, stability and YAP target gene expression, we wanted to examine if JM7 inhibits cell proliferation, colony formation and migration of MDA-MB-231 and OVCAR-8 ovarian cancer cells that exhibit high YAP activity. To test the effect of JM7 on cell proliferation, we treated MDA-MB-231 and OVCAR-8 cells with either DMSO or 2 micromolar JM7 and performed the MTT assay. In this assay, the colorless MTT reagent is converted by the cellular oxidoreductases to colored formazan crystals, which are then solubilized and quantitated by measuring the absorbance of the colored product. Thus, it provides an indirect measure of the number of cells. We observed that JM7 treatment significantly impacts the proliferation of these cancer cells (Figure 6. A, B). We then examined if JM7 affects the colony forming ability of these cells. To address this, we plated equal number of MDA-MB-231 and OVCAR8 cells and treated them with DMMSO or 1, 2 or 5 micromolar JM7 for 2 weeks, after which the cells were fixed and stained with crystal violet. We observed that even at 1um dose JM7 impacted colony forming ability of the MDA-MB-231 cells. Interestingly, at 2 and 5 micromolar there was a dramatic reduction in the colony forming ability of these cells (Figure 6. C, E). On the other hand, JM7 had a significant effect on the colony formation ability of OVCAR-8 at 5 micrommolar (Figure 6. D, Fs). Finally, we wanted to examine how JM7 affects the migration of the MDA-MB-231 and OVCAR8 cells. To address this, we grew the MDA-MB-231 and OVCAR8 cells in confluent monolayers and scratched with a pipette tip. After carefully washing away the detached cells, we treated them with DMSO alone or 2 micromolar JM7 and imaged 0, 19 and 24 hours after drug treatment. We observed that while cells treated with DMSO alone gradually migrated and filled the gap over time, JM7 treatment significantly attenuated this response (Figure 6. G-L). Together, these experiments indicate that JM7 inhibits proliferation, colony forming ability and migration of the MDA-MB-231 and OVCAR-8 cancer cells.

**Figure 6.**
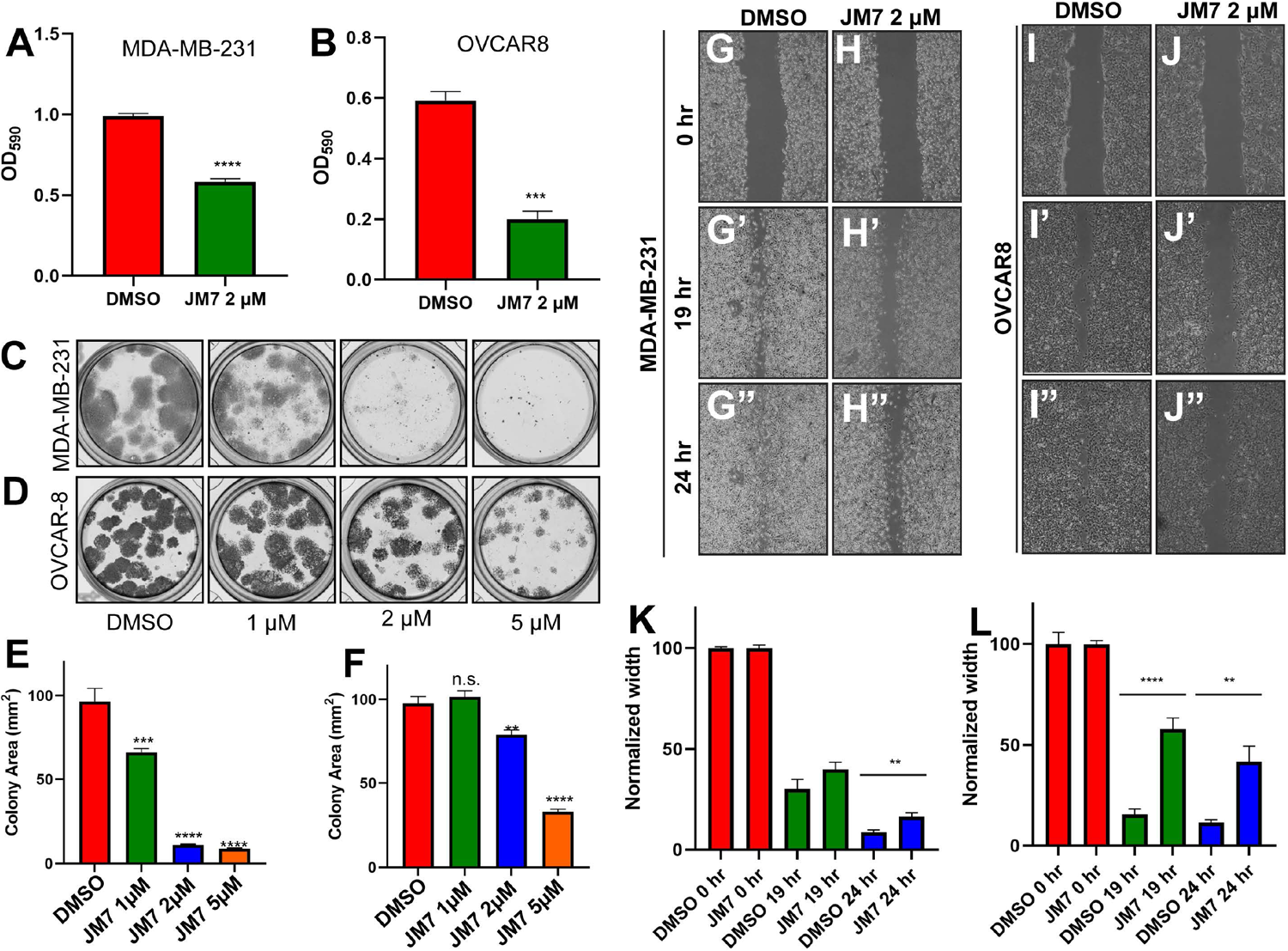
JM7 inhibits proliferation, colony formation and migration of breast and ovarian cancer cells. (A,B). MTT assay for MDA-MB-231 cells (A) and OVCAR-8 cells (B) treated with DMSO or the indicated doses of JM7, showing inhibition of proliferation of these cancer cells by JM7. (C-F”) Representative images showing colony formation assay for MDA-MB-231 (C) and OVCAR-8 cells (D) treated with DMSO or 1, 2 or 5 micromolar JM7. (E, F) Histograms showing normalized colony area for DMSO and JM7 treated MDA-MB-231 (E) and OVCAR-8 (F). (G-J”) Representative images showing wound healing assay for MDA-MB-231 (G-H”) and OVCAR-8 cells (I-J”) treated with DMSO or 2 micromolar JM7. (I, J) Histograms showing normalized wound width for DMSO and JM7 treated MDA-MB-231 (I) and OVCAR-8 (J) cells at 0, 19 and 24 hours. ** p <0.05; *** p <0.005; **** p <0.001. Error bars indicate Standard Error of Mean (SEM). Please also see Figure S2.

## 4 DISCUSSION

The transcriptional coactivators YAP/TAZ are the distal effectors of the Hippo signaling pathway and play a critical role in regulating cell proliferation, apoptosis and fate specification (1, 2). They regulate expression of majority their target genes by interacting with TEAD1-4 transcription factors and are frequently deregulated in several cancers, where they regulate multiple aspects of cancer development including cancer growth, metastasis and chemo/immunotherapy resistance (8). Thus, they provide a critical target for therapeutic intervention. However, it is not possible to directly target YAP/TAZ since they are intrinsically disordered. Therefore, YAP activity can be alternatively controlled by targeting TEAD1-4, which contain a highly druggable central hydrophobic pocket that is normally occupied by a palmitic acid. In an effort to develop a small molecule inhibitor for TEAD, in this study we undertook a structure-based computational screening of a large library of small-molecule inhibitors and identified a compound that inhibits YAP transcriptional activity. It directly binds to TEAD in cells and inhibits their palmitoylation and decreases their stability. Further, it inhibits proliferation, colony formation and migration of MDA-MB-231 and OVCAR-8 cancer cells.

It has been shown that TEAD palmitoylation is not required for its nuclear localization but required for TEAD stability and its interaction with YAP/TAZ (31). Consistent with this, we observed that JM7 inhibited palmitoylation of all four TEAD isoforms, but did not affect TEAD or YAP localization. Further, consistent with previous reports, we observed that JM7 treatment destabilized all the 4 TEAD isoforms. Interestingly, we observed JM7 affects stability of the different TEAD isoforms differently. Especially, the destabilizing effect was not as drastic in case of TEAD3. This could be due to less efficient inhibition of palmitoylation or higher expression of TEAD3. Another possibility could be that the different TEAD isoforms have different stability in unpalmitoylated state. Nevertheless, isoform specific TEAD-inhibitors are valuable tools to studying TEAD biology and as potential anticancer agents with selectivity and less toxicity. Further structure activity studies will be required to develop analogs that can potently inhibit all the four isoforms.

Loss of NF2 function is commonly detected in many cancers including malignant mesothelioma, meningioma and schwannoma (10). MDA-MB-231 cells also harbor deletions in NF2, which causes activation of YAP in these cells (41). JM7 inhibits YAP target gene expression, viability, proliferation and migration of these cells. Further investigations will be required to examine if JM7 is effective in other cancers with NF2 mutation. Similarly, in many cancers YAP/TAZ are overexpressed by upstream oncogenic signaling pathways such as EGFR, RAS-MAPK, PI3, WNT, and exhibit increased nuclear localization. Therefore, we posit that JM7 will be effective in many cancers that exhibit high expression and nuclear localization of TAP/TAZ. Further increased YAP nuclear localization has also been reported to be associated with chemotherapy resistance and relapse in EGFR-mutant, KRAS-mutant and B-RRAF mutant, ALK-rearranged non-small-cell lung cancer and RAS-driven neuroblastoma (19, 22, 42-48). TEAD palmitoylation inhibitors such as JM7 will be useful adjunct therapy along with these pathway specific inhibitors. Similarly, increased YAP activity in cancer cells and immune cells interferes with immunotherapy (19, 22, 24-28). Therefore, JM7 will be useful in combination with immunotherapy such as immune check point inhibitors.

YAP and TAZ also play a critical role in cancer associated fibroblasts and stimulate fibrosis, which in turn activates YAP, thereby creating a vicious feed forward loop. Similarly, they play a critical role in pulmonary, hepatic and renal fibrosis, where downstream of TGF beta signaling, YAP/TAZ play a pivotal role in promoting conversion of the fibroblasts into myofibroblasts and induce expression of genes encoding ECM components (49). We propose that JM7 may be effective in fibrotic conditions, where YAP/TAZ plays a prominent role. Further investigations will be required for assessing the in vivo efficacy of JM7 in cancer and fibrosis. Moreover, new analogs can be synthesized and tested to develop more potent TEAD inhibitors with desirable pharmacological properties.

## Acknowledgments

We thank the Misra lab members for their critical inputs on the manuscript. We thank Nikki Delk for the kind gift of MDA-MB-231 cells and Dr. Girgis Obaid for kindly allowing us to use the Gelcount colony imager. This research was funded by University of Texas at Dallas Start-up funds to JRM.

## Author Contributions

Conceptualization, JRM; methodology, JRM.; validation, AG, SM, JRM.; formal analysis, JRM; investigation, AG, SM, JRM; resources, JRM.; data curation, JRM writing-original draft preparation, JRM; writing-review and editing, JRM.; visualization, AG, SM, JRM supervision, JRM; project administration, JRM; funding acquisition, JRM. All authors have read and agreed to the published version of the manuscript.

### Data Availability statement

The data that support the findings of this study are available in the methods and/or supplementary material of this article. Data sharing not applicable to this article as no large datasets were generated or analyzed during the current study

## SUPPLEMENTAL INFORMATION

**Supplemental Figures S1 and S2**

**Supplemental Table-1**

## SUPPLEMENTAL FIGURES

**Figure S1. YAP5S and TAZS89A activates the 8XGTIIC reporter**.

A) Histograms showing relative Fluc activity in HEK293 stable cells carrying 8XGTIIC-Fluc reporter transfected with empty vectors or plasmids encoding YAP5SA (A) or TAZS89A (B), indicating very low basal activity of the reporter and hyperactivation of the reporter by YAP5SA and TAZS89A. Error bars indicate standard error of mean (SEM)

**Figure S2. Validation of the Nanobit complementation assay**.

A-D) Histograms showing relative Nanoluc activity in HEK293 SmBit-Yap (E-H) or Smbit-TAZ (I-L) together with TEAD1(A), TEAD2 (B), TEAD3 (C) and TEAD4 (D). Error bars indicate Standard Error of Mean.

**Table S1**

List of key resources used in this study.

**Table S2**

List of oligonucleotides used for molecular cloning in this study.

