## Supplemental information for "Structure-based discovery of a novel small-molecule inhibitor of TEAD palmitoylation with anticancer activity"

**Supporting information**

**Authors:** Artem Gridnev, Subhajit Maity, Jyoti R. Misra

Department of Biological Sciences

University of Texas at Dallas, Richardson, TX, United States

**Index**

**Supplemental Figure-1:** YAP5SA and TAZS89A hyperactivates the 8XGTIIC reporter……………2
**Supplemental Figure-1 :** Validation of the nanobit complementation assay…………………………..3
**Supplemental Table-1:** List of key resources used in this study………………………………………4
**Supplemental Table-2:** List of oligonucleotides used for molecular cloning in this study…………….6

*Supplementary figures*

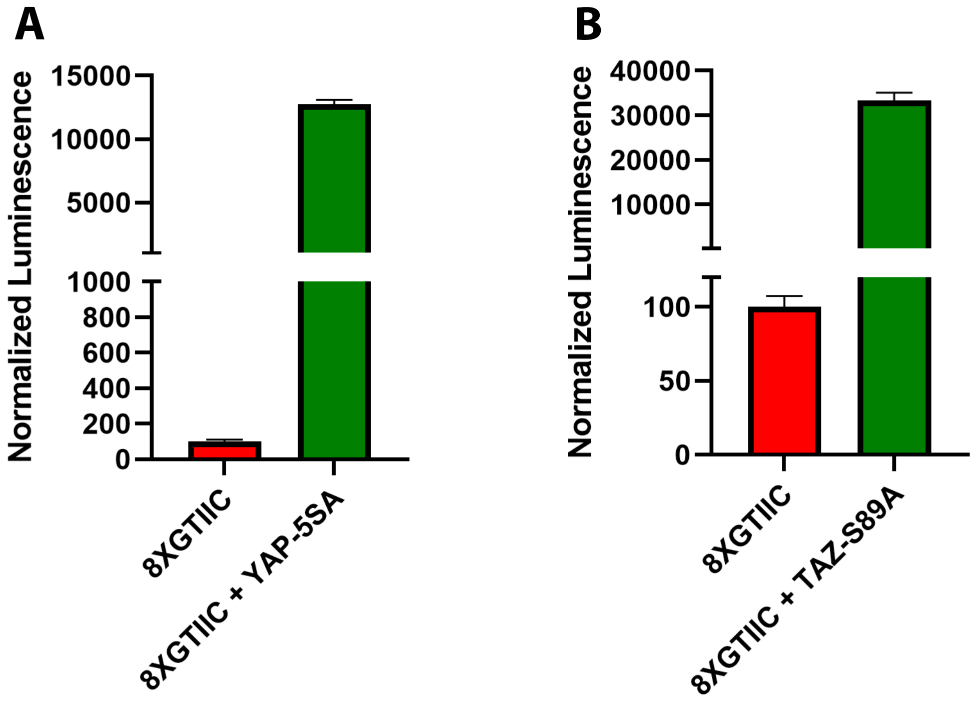


**Figure S1. YAP5SA and TAZS89A hyperactivates the 8XGTIIC reporter.**
A) Histograms showing relative Fluc activity in HEK293 stable cells carrying 8XGTIIC-Fluc reporter transfected with empty vectors or plasmids encoding YAP5SA (A) or TAZS89A. (B), indicating very low basal activity of the reporter and hyperactivation of the reporter by YAP5SA and TAZS89A. Error bars indicate standard error of mean (SEM)

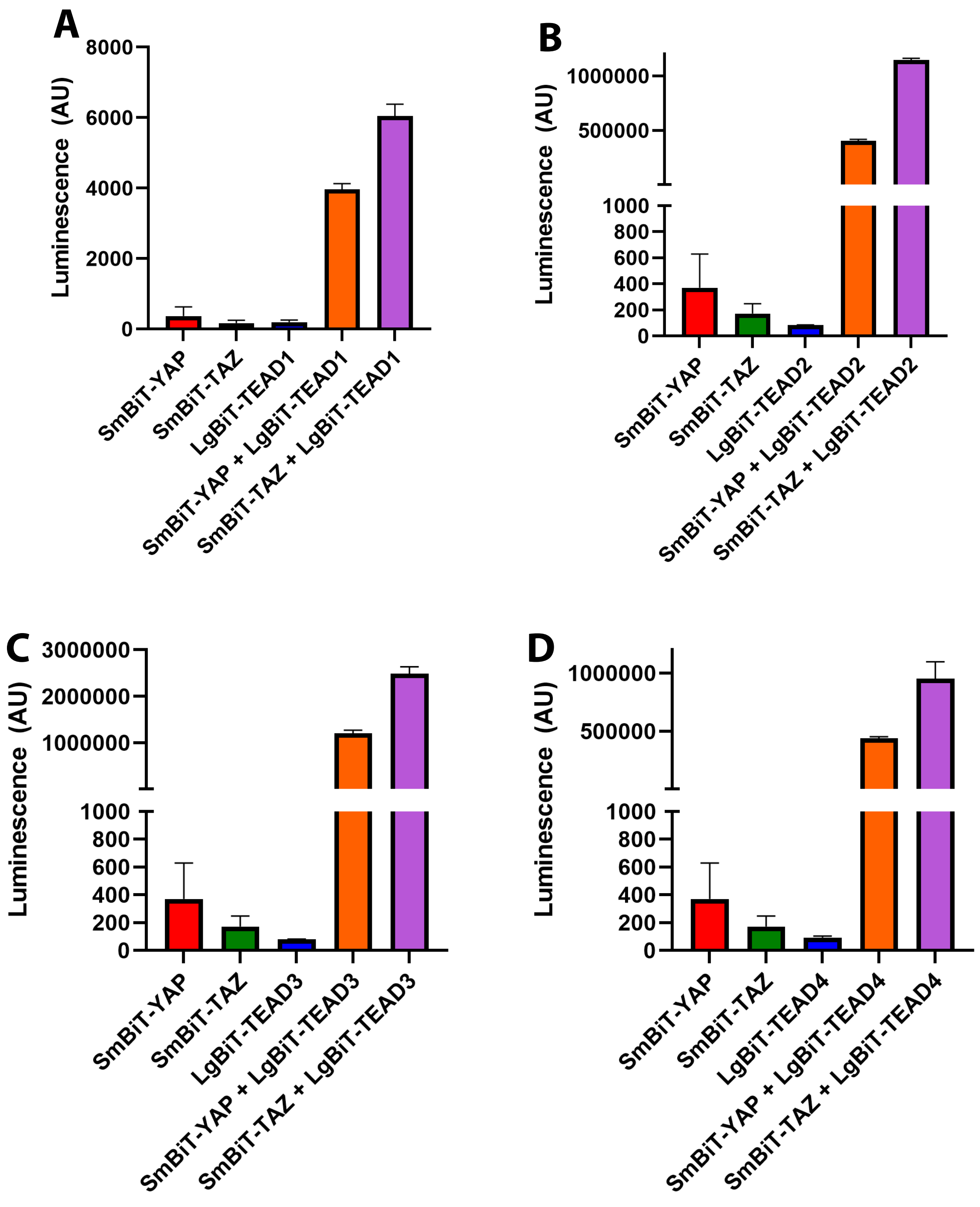


**Figure S2. Validation of the nanobit complementation assay.**
A-D) Histograms showing relative Nanoluc activity in HEK293 SmBit-Yap (E-H) or Smbit-TAZ. (I-L) together with TEAD1(A), TEAD2 (B), TEAD3 (C) and TEAD4 (D). Error bars indicate Standard Error of Mean.

**Supplementary Table-1**

List of key resources used in this study.

| REAGENT or RESOURCE | SOURCE | IDENTIFIER |
|---|---|---|
| **Antibodies** | | |
| YAP (D8H1X) Rabbit mAb | Cell Signaling Technology | 14074 |
| Pan-TEAD (D3F7L) Rabbit mAb | Cell Signaling Technology | 13295 |
| Myc-Tag (9B11) Mouse mAb | Cell Signaling Technology | 2276 |
| Myc-Tag (71D10) Rabbit mAb | Cell Signaling Technology | 2278 |
| Pierce Anti c-Myc Agarose | Fisher Scientific | PI20168 |
| Goat anti-Rabbit IgG IRDye 800CW conjugate | LI-COR | 925-32211 |
| Goat anti-Mouse IgG IRDye 680RD conjugate | LI-COR | 925-68070 |
| Alexa Fluor® 647 AffiniPure Donkey Anti-Rabbit IgG (H+L) | Jackson ImmunoResearch | 711-605-152 |
| Bacterial and virus strains | | |
| Mach1 T1$^R$ Chemically Competent *E. coli* | ThermoFisher | C862003 |
| TEAD Luciferase Reporter Lentivirus | BPS Bioscience | 79833 |
| **Chemicals, peptides, and recombinant proteins** | | |
| DyLight 680-Conjugated Streptavidin | Rockland Immunochemicals | S000-44 |
| JM7 | Vitas-M Laboratory | STK096693 |
| Alkynyl Palmitic Acid | Click Chemistry Tools | 1165 |
| Biotin Azide | Click Chemistry Tools | 1265 |
| tris (3-hydroxypropyltriazolylmethyl) amine (THPTA) | Click Chemistry Tools | 1010 |
| Tris(2-carboxyethyl) phosphine (TCEP) hydrochloride solution | Sigma-Aldrich | 646547 |
| Copper (II) sulfate pentahydrate | Sigma-Aldrich | C8027 |
| IGEPAL-CA630 | Sigma-Aldrich | I3021 |
| Invitrogen TRIzol Reagent | ThermoFisher | 15596026 |
| Applied Biosystems Fast SYBR Green Master Mix | ThermoFisher | 4385612 |
| Lipofectamine 3000 | ThermoFisher | L3000015 |
| Hoechst 33342 | ThermoFisher | H3570 |
| Vectashield mounting medium | Vector Laboratories | H1000 |
| **Critical commercial assays** | | |
| Dual-Luciferase Reporter Assay System | Promega | E1980 |
| Nano-Glo Luciferase Assay System | Promega | N1120 |
| CellTiter-Glo 2.0 Cell Viability Assay | Promega | G9242 |
| ONE-Step Luciferase Assay System | BPS Bioscience | 60690 |
| MTT Assay Kit | abcam | ab211091 |
| NEBuilder HiFi DNA Assembly Master Mix | New England BioLabs | E2621 |
| **Experimental models: Cell lines** | | |
| HEK293 | ATCC | CRL-1573 |
| MDA-MB-231 | ATCC | HTB-26 |

| | | |
|---|---|---|
| OVCAR8 | NCI | |
| **Oligonucleotides** | | |
| Please refer to table S1 | | |
| **Recombinant DNA** | | |
| pCMX-GAL4-TEAD1 | Addgene | 33108 |
| pCMX-GAL4-TEAD2 | Addgene | 33107 |
| pCMX-GAL4-TEAD3 | Addgene | 33106 |
| pCMX-GAL4-TEAD4 | Addgene | 33105 |
| pRK5-Myc-TEAD4 | Addgene | 24638 |
| pCMV-Flag-YAP-5SA/S94A | Addgene | 33103 |
| GST-YAP2 | Addgene | 24637 |
| pCDNA3-HA-TAZ | Addgene | 32839 |
| pRL-TK | Promega | E2241 |
| Myc-TEAD3 gBlock | IDT | |
| pDONR221-TEAD3 | DNASU | HsCD00963910 |
| pCDNA3.1-TAZ-S89A | This study | |
| pCDNA3.1-Myc-TEAD1 | This study | |
| pCDNA3.1-Myc-TEAD2 | This study | |
| pCDNA3.1-Myc-TEAD3 | This study | |
| pCDNA3.1-Myc-TEAD4 | This study | |
| pCDNA3.1-SmBiT-YAP-FLAG | This study | |
| pCDNA3.1-SmBiT-TAZ-FLAG | This study | |
| pCDNA3.1-LgBiT-TEAD1-myc | This study | |
| pCDNA3.1-LgBiT-TEAD2-myc | This study | |
| pCDNA3.1-LgBiT-TEAD3-myc | This study | |
| pCDNA3.1-LgBiT-TEAD4-myc | This study | |
| **Software and algorithms** | | |
| Fiji | NIH | https://imagej.net/Fiji; RRID: SCR_002285 |
| Schrödinger software | Schrödinger Inc | https://www.schrodinger.com/ |
| Prism 9 | GraphPad | https://www.graphpad.com/ |
| Volocity | PerkinElmer | https://www.perkinelmer.com/ |
| LAS X Life Science Microscope Software | Leica | https://www.leica-microsystems.com/products/microscope-software/p/leica-las-x-ls/ |

**Supplemental Table1**

**List of oligonucleotides used for molecular cloning in this study.**

| Oligo name | Sequence (5'-3') |
|---|---|
| pCDNA3.1-myc-TEAD1 FWD | TAGTCCAGTGTGGTGGAATTCGCCACCATGGAGCAGAAGCTGATCAGCGAGGAGGACCTGATGAGTGACTCTGCAGATAAGCCA |
| pCDNA3.1 TEAD1 REV | TGCTGGATATCTGCAGAATTCCTAGTCCTTTACAAGCCTGTAAATATGATGTTGTG |
| pCDNA3.1-myc-TEAD2 FWD | TAGTCCAGTGTGGTGGAATTCGCCACCATGGAGCAGAAGCTGATCAGCGAGGAGGACCTGGGGGAACCCCGGGCT |
| pCDNA3.1 TEAD2 REV | TGCTGGATATCTGCAGAATTCCTAGTCCCTGACCAGGCGG |
| pcDNA3.1 LgBiT FWD | TAGTCCAGTGTGGTGGAATTCGCCACCATGGTCTTCACACTCGAAGATTTCGT |
| pCDNA3.1 LgBiT TEAD1 REV | TGCTGGATATCTGCAGAATTCCTACAGATCCTCTTCTGAGATGAGTTTTTGTTCA |
| LgBiT REV TEAD2 OH | TGGGGGCAGGGGTGAACCGCTCGAGCCTCC |
| TEAD2 FWD LgBiT OH | GGAGGCTCGAGCGGTTCACCCCTGCCCCCAC |
| pCDNA3.1 LgBiT TEAD2 REV | TGCTGGATATCTGCAGAATTCCTACAGATCCTCTTCTGAGATGAGTTTTTGTTCGTCCCTGACCAGGCGG |
| LgBiT REV TEAD3 OH | ACTAGGGAGTGGTGCACCGCTCGAGCCTCC |
| TEAD3 FWD LgBiT OH | GGAGGCTCGAGCGGTGCACCACTCCCTAGTGCC |
| pCDNA3.1 LgBiT TEAD3 REV | TGCTGGATATCTGCAGAATTCCTACAGATCCTCTTCTGAGATGAGTTTTTGTTCATCTTTCACCAGCTTGTACACGT |
| LgBiT REV TEAD4 OH | GGGCGATGGGGCGGGACCGCTCGAGCCTCC |
| TEAD4 FWD LgBiT OH | GGAGGCTCGAGCGGTCCCGCCCCATCGC |
| pCDNA3.1 LgBiT TEAD4 REV | TGCTGGATATCTGCAGAATTCCTACAGATCCTCTTCTGAGATGAGTTTTTGTTCTTCTTTCACCAGCCTGTAGATGTGG |
| pCDNA3.1 SmBiT FWD | TAGTCCAGTGTGGTGGAATTCGCCACCATGGTGACCGGCTACCG |
| pCDNA3.1 SmBiT YAP REV | TGCTGGATATCTGCAGAATTCCTACTTGTCGTCATCGTCTTTGTAGTCTACAT |
| pCDNA3.1 SmBiT TAZ REV | TGCTGGATATCTGCAGAATTCCTACTTGTCGTCATCGTCTTTGTAGTCGT |
| pCDNA3.1 TAZ FWD | TAGTCCAGTGTGGTGGAATTCATGGCCTACCCATACGATGTTCC |
| TAZ S89A REV | CAGGGACGCGGGCGAGGCGTGCGAGCGGACATGTTGG |
| TAZ S89A FWD | CATGTCCGCTCGCACGCCTCGCCCGCGTCCCT |
| pCDNA3.1 TAZ REV | TGCTGGATATCTGCAGAATTCTTACAGCCAGGTTAGAAAGGGCTC |
| GAPDH FWD qPCR | GAAGGTCGGAGTCAACGGATT |
| GAPDH REV qPCR | CGCTCCTGGAAGATGGTGAT |
| CTGF FWD qPCR | GTTTGGCCCAGACCCAACTA |
| CTGF REV qPCR | GGCTCTGCTTCTCTAGCCTG |
| CYR61 FWD qPCR | CAGGACTGTGAAGATGCGGT |
| CYR61 REV qPCR | GCCTGTAGAAGGGAAACGCT |
